# A brain model of altered self-appraisal in social anxiety disorder

**DOI:** 10.1101/2023.04.26.537105

**Authors:** Alec J. Jamieson, Ben J. Harrison, Rebekah Delahoy, Lianne Schmaal, Kim L. Felmingham, Lisa Phillips, Christopher G. Davey

## Abstract

The brain’s default mode network has a central role in the processing of information concerning oneself. Dysfunction in this self-referential processing represents a key component of multiple mental health conditions, including social anxiety disorder (SAD). This case-control study aimed to clarify alterations to network dynamics present during self-appraisal in SAD participants. A total of 38 adolescents and young adults with SAD and 72 healthy control participants underwent a self-referential processing fMRI task. The task involved two primary conditions of interest: *direct self-appraisal* (thinking about oneself) and *reflected self-appraisal* (thinking about how others might think about oneself). Dynamic causal modelling and parametric empirical Bayes were then used to explore differences in the effective connectivity of the default mode network between groups. We observed connectivity differences between SAD and healthy control participants in the reflected self-appraisal but not the direct self-appraisal condition. Specifically, SAD participants exhibited greater excitatory connectivity from the posterior cingulate cortex (PCC) to medial prefrontal cortex (MPFC) and greater inhibitory connectivity from the inferior parietal lobule (IPL) to MPFC. In contrast, in the absence of task modulation, SAD participants exhibited reduced intrinsic connectivity, with reduced excitatory connectivity from the PCC to MPFC and reduced inhibitory connectivity from the IPL to MPFC. As such, participants with SAD showed changes to afferent connections to the MPFC which occurred during both reflected self-appraisal as well as intrinsically. The presence of connectivity differences in reflected and not direct self-appraisal is consistent with the characteristic fear of negative social evaluation that is experienced by people with SAD.

## Introduction

Social anxiety disorder (SAD) is one of the most common mental illnesses, with a lifetime prevalence of 11% ^1^. Adolescence is frequently identified as the primary period for the development of SAD, with a median age of onset in early adolescence^2, 3^ and a peak prevalence in later adolescence^4^. People with SAD often experience significant distress during social encounters^5^, and in turn, make considerable efforts to avoid them. As a result of these factors, SAD has a dramatic effect on social and educational functioning throughout a critical period of development^6^ and often results in reduced quality of life^7^. Despite this, only a minority of people with anxiety disorders receive adequate treatment^8^. Current guidelines emphasize the use of selective serotonin and noradrenalin reuptake inhibitors, in addition to psychological treatments including cognitive behavioral therapy (CBT) as first-line interventions^9, 10^. Notably, both psychological and pharmacological treatments have demonstrated small to moderate effect sizes for those with anxiety disorders^11^.

An essential characteristic of SAD is a consistent fear of negative evaluation, which can result in excessively focusing on themselves during social interactions. Their self-representations are influenced by how they imagine their appearance and behavior are perceived by others, with fears that they are viewed as socially inept, gauche, and awkward^12^. This disturbance in self-appraisal processes is associated with marked anxiety, and is one of the features of SAD that is targeted by CBT^13^.

Functional MRI studies of participants with SAD often show alterations in brain regions that support self-appraisal processes. The medial prefrontal cortex (MPFC) and posterior cingulate cortex (PCC) are core regions of the default mode network (DMN), a collection of brain regions which are highly active during self-appraisal^14, 15^. SAD associated alterations have been observed in both of these regions, with the MPFC showing increased activation during self-appraisal processes^16, 17^ and the PCC showing increased activation in response to facial stimuli^18, 19^. Those with SAD have also been shown to demonstrate increased recruitment across DMN regions during reappraising compared with accepting negative self-beliefs^20^. Similarly, altered functional connectivity between the MPFC and PCC has been illustrated in SAD participants^21-24^, although the directionality of these effects appears inconsistent. Alterations between DMN regions have also been observed in other anxiety disorders^25-27^, as well as being associated with trait^28^ and state anxiety levels^29, 30^. Together this evidence indicates that alterations to the DMN likely generalize beyond SAD.

In addition to these midline regions, self-appraisal is associated with increased activity and connectivity with a third component of the DMN, the inferior parietal lobule (IPL). Using effective connectivity (the directional effect that activity in one region has on the activity of another), we have previously shown that the PCC exerts a positive influence on the IPL and MPFC during self-appraisal, with the latter in turn exerting negative feedback on the PCC^15^. In a more recent study, we compared direct self-appraisal (where the participant assessed whether personality adjectives described them) with reflected self-appraisal (where the participant assessed whether *other* people would think personality adjectives described them) in healthy participants^31^. The latter task component showed similar patterns of activity and directed connectivity, although with greater activation of the PCC and IPL, and a distinct pattern of negative influence of the IPL on PCC.

The examination of reflected self-appraisal is likely to be particularly pertinent for people with SAD, considering the marked anxiety associated with their fear of negative appraisal by others inherent to the condition. Our aim was to examine brain connectivity in participants with SAD as they engaged in a task that examined direct and reflected self-appraisal processes. As in our previous studies, we aimed to use dynamic causal modelling to examine effective connectivity, comparing participants with SAD with control participants across the two conditions. We hypothesized that connectivity differences would be more apparent for SAD participants performing reflected self-appraisal than direct self-appraisal. Consistent with our previous work^32^, we propose that these differences will likely occur in the modulation from the MPFC to PCC.

## Methods and Materials

### Participants

Forty-two unmedicated SAD participants were recruited from outpatient mental health clinics in the western and northern suburbs of Melbourne, Australia. These participants had a primary diagnosis of SAD, as assessed with the Structured Clinical Interview for DSM-5 Axis I disorders (SCID-5-RV)^33^. SAD participants’ symptoms were in the moderate to severe range on the Liebowitz Social Anxiety Scale (LSAS)^34^. Imaging data from 4 SAD participants were excluded due to excessive head motion (N = 1), and suboptimal performance of the external attention component of the task (N = 3; see below), leaving a total sample of 38 SAD participants.

We recruited 122 healthy controls with no current or past diagnoses of mental illness (assessed with the SCID-5 non-patient interview)^33^, which we have reported on previously^31^. Exclusion due to excessive head motion (N = 3), inadequate task performance (N = 8), and insufficient subject level activation for DCM (N = 7), resulted in 104 participants with adequate data. To ensure that participants did not differ in key covariates, we conducted variable ratio matching using age and gender with the *MatchIt* package in R^35^ (see Supplementary Materials for full details). This resulted in a subsample of 72 matched controls being included in these analyses.

All participants met the following inclusion criteria: they were aged between 16 and 25 years, were competent English speakers, had no current treatment with psychoactive medications, had no dependence on alcohol or other drugs, showed no incidental neurological findings on MRI, and had no further contraindications to MRI. Written informed consent was obtained from each participant following a complete description of the study protocol. For participants under 18 years, both participant assent and parent consent were obtained. The study protocol was approved by The University of Melbourne Human Research Ethics Committee and was carried out in accordance with the Declaration of Helsinki.

### Task design

Participants completed an fMRI task composed of 4 experimental conditions: direct self-appraisal, reflected self-appraisal, external attention, and rest-fixation – the same task design described by Delahoy and colleagues^31^. In 3 of the conditions, participants were presented with words drawn from a list of frequently used trait adjectives^36^. To heighten self-reflection, words were selected to be neither extremely favorable nor unfavorable. As such, selected adjectives were from the subset of words rated as most ‘meaningful’ and were distributed around the median rating for ‘likeableness’. Words were presented in blocks (6 blocks per condition, 5 words per block) and the word order was randomized for each participant. In the direct self-appraisal condition, they were asked to respond to the question “Would you use this word to describe you?”. In the reflected self-appraisal condition, they were asked “Would others use this word to describe you?”, and in the external attention condition they responded to the question “Does this word have four or more vowels?”. To each stimulus, they responded “Yes” or “No” by pressing one of two allocated buttons on a curved 4-button fiber-optic response pad (Cambridge Research Systems Ltd.). The 3 lists of 30 words that formed these conditions were matched on valence and number of vowels and were counterbalanced across participants (Supplementary Table S1). Each 27-second block (2 seconds of instruction followed by 5 words presented for 5 seconds each) was interspersed with a 10-second rest-fixation block in which participants were asked to fixate on a centrally presented crosshair.

### Image acquisition

A 3-T General Electric Discovery MR750 system equipped with an 8-channel phased-array head coil was used in combination with ASSET parallel imaging. Blood oxygenation level-dependent imaging was undertaken using a single-shot gradient-recalled echo planar imaging sequence in the steady state (repetition time, 2000 ms; echo time, 33 ms; pulse angle, 90º) in a 23 cm field of view, with a 64x64 pixel matrix and a slice thickness of 3.5 mm (no gap). Thirty-six interleaved slices were acquired parallel to the anterior-posterior commissure line with a 20º anterior tilt to better cover ventral prefrontal cortical regions. The total sequence time was 11 min 24 s, corresponding to 342 whole-brain echo-planar imaging volumes. A T1-weighted high-resolution anatomical image was acquired for each participant to assist with functional time-series co-registration (140 contiguous slices; repetition time, 8.2 ms; echo time, 3.2 ms; flip angle, 13º; in a 25.6 cm field-of-view, with a 256x256 pixel matrix and a slice thickness of 1 mm). To assist with noise reduction and head immobility, participants were fitted with insert-ear protection and their heads were supported with foam-padding inserts.

### Image preprocessing

Imaging data were transferred to a Unix-based platform that ran MATLAB version 9.3 (The MathWorks Inc., Natick, USA) and Statistical Parametric Mapping Version 12 v7771 (SPM12; Wellcome Trust Centre for Neuroimaging, UK). Each participant’s time-series were aligned to the first image using least-squares minimization and a 6-parameter rigid-body spatial transformation. Motion Fingerprint toolbox (Wilke, 2012) was used to characterize scan-to-scan head motion for each participant. Participants’ data were excluded if scan-to-scan displacement exceeded 2.5 mm or total displacement exceeded 3 mm (∼1 native voxel). Realigned and slice time corrected functional images were then co-registered to each participant’s respective T1 anatomical scans, which were segmented and spatially normalized to the International Consortium of Brain Mapping template using the unified segmentation approach. The functional images were transformed to 2 mm isotropic resolution and were smoothed with an 8 mm full-width-at-half-maximum Gaussian filter.

### General linear modelling

Following preprocessing, first-level general linear model (GLM) analyses were conducted for each participant in SPM12. To predict the activity of a given voxel as a function of time, regressors of interest were generated by specifying the durations and onsets of each block for the direct self-appraisal, reflected self-appraisal, rest-fixation, external attention, and instruction conditions. These were then convolved with a canonical hemodynamic response function. The external attention condition was considered an appropriate baseline as it was matched with the self-appraisal conditions on stimulus features yet required specific attentional demands to minimize the intrusion of task-independent thought. A high-pass filter (1/128 s) accounted for low-frequency drifts. GLM with local autocorrelation correction was used to calculate parameter estimates at each voxel. Primary contrast images from each participant were carried forward to second-level analyses using the summary statistics approach to random-effects analyses. Independent sample *t*-tests were used for between-groups analyses, with a whole brain threshold family-wise error rate (FWE) corrected threshold of *P* < 0.05, K_E_ ≥ 20 voxels.

Following Delahoy and colleagues^31^, we then applied conjunction (null) analyses to identify “core-self DMN” regions across groups: those that were commonly activated by both self-appraisal conditions as well as rest-fixation, but that showed further distinct activation during the self-appraisal conditions relative to rest-fixation. These core-self regions were used as volumes of interest (VOIs) for further analysis with DCM. Results are reported in Montreal Neurological Institute space.

### Dynamic causal modelling

DCM is a method for estimating the directional interactions between brain regions from observed neuroimaging data^37^. For tasks, this estimation is demarcated into invariant connectivity in the absence of task modulation (intrinsic connectivity) and connectivity that is modulated by experimental stimuli^38^. For interactions between regions, the strength of this relationship is measured in hertz (Hz) and positive values indicate excitation while negative values indicate inhibition. In contrast, self-connections are unitless log scaling parameters for which positive values indicate greater inhibition whereas negative values indicate reduced inhibition.

As DCM is a hypothesis-driven method requiring selection of *a priori* regions of interest, we restricted our analysis to “core-self DMN regions” as defined in our previous studies^15, 31^. These regions included the MPFC, PCC, and left IPL and were selected due to their strong association with self-related processing^39-41^. To localize and identify peak coordinates of these regions, we used the aforementioned conjunction analysis. The time-series for our chosen VOIs were then extracted at a single subject level using the first eigenvariate of voxels within 5 mm of the subject-specific maxima that showed significant activation for the rest-fixation and direct and reflected self-appraisal > external attention contrast (*P* < 0.05). If a given VOI had inadequate activation, the threshold was incrementally relaxed up to *P* < 0.5^38^. Specific coordinates for all three regions were personalized to each subject’s specific local maximum, which were required to occur within 8 mm of the group level peaks identified in the conjunction analysis (see Supplementary Figure S1). A full model was specified with intrinsic bidirectional connections between the MPFC, PCC, and left IPL (Supplementary Figure S2) and modulation for both the direct and reflected self-appraisal conditions occurring for all connections. Direct input into the network was modeled using the overall ‘broad self’ (direct + reflected + rest) into the PCC and the input matrix was not mean centered. As such, the intrinsic connectivity represents the connectivity when there is driving input present and task modulation is absent (e.g., connectivity during the rest fixation). Full models were estimated separately for each subject.

We deployed Parametric Empirical Bayes (PEB) to examine the differences in connectivity parameters between healthy controls and SAD participants^42^. PEB enables the inclusion of the estimated variance of connectivity parameters when investigating between-group effects, thereby allowing for more reliable parameter estimates than using classical tests. While Bayesian statistics has no concept of statistical significancy, thresholding can be used to identify effects which are likely of interest. We used a posterior probability (PP) threshold of greater than 0.95^43^. The regressors included in this analysis represented 1) the average connectivity across all participants, 2) the effect of SAD, and 3) the effect of age. An automatic search over nested PEB models was conducted following the estimation of a group level PEB model^44, 45^. After the final iteration, Bayesian model averaging was performed to determine the strength of connections in the last Occam’s window of 256 models.

### Leave-one-out cross-validation

To assess the predictive validity of parameters which demonstrated between group differences we used leave-one-out cross-validation (LOOCV) across both groups. In the PEB framework, LOOCV aims to determine whether the size of between-group effects on parameters is sufficiently large to predict a variable of interest^43^, in this case, anxiety symptom severity. This does so by estimating a group-level PEB model while excluding one subject, then using this PEB model to predict the left-out subject’s anxiety symptom severity. This predicted anxiety score is then correlated with the observed anxiety symptom severity. A significant correlation between the expected and observed values demonstrates that the effect size was sufficiently large to predict the left-out subjects’ anxiety symptom severity above chance (see^43^ for further details). Anxiety symptom severity was assessed through the LSAS (social anxiety symptoms), as well as the State-Trait Anxiety Inventory (STAI)^46^ general and present subscales (trait and state anxiety symptoms respectively).

## Results

### Clinical and demographic characteristics

Comparison of clinical characteristics revealed that SAD participants had significantly higher LSAS (*t*(56.26) = -13.65, *P* < .001), STAI General (*t*(108) = -13.52, *P* < .001) and STAI Present scores (*t*(108) =-9.49, *P* < .001) compared with healthy controls (Table 1). For distribution of anxiety severity scores across groups see Supplementary Figure S3. No significant differences were observed between groups for age, gender, or IQ (Table 1).

**Table 1.** Demographic and Behavioral Differences Between Healthy Controls and Social Anxiety Disorder Participants.

| Characteristics | Healthy Controls<br>(*N* = 72) |  | SAD participants<br>(*N* = 38) |  | Cramer's *V*<br>or Hedges' *g* | *P* |
| --- | --- | --- | --- | --- | --- | --- |
|  | *N* or<br>Mean | Percentage<br>or *SD* | *N* or<br>Mean | Percentage<br>or *SD* |  |  |
| Female | 45 | 62.5 | 23 | 60.5 | .03 | .826 |
| Comorbid MDD | - | - | 25 | 65.8 | - | - |
| Age (years) | 20.5 | 1.8 | 19.8 | 2.3 | .35 | .166 |
| Standardized IQ | 111.9 | 8.4 | 110.3 | 13.4 | .15 | .816 |
| Liebowitz Social Anxiety<br>Scale score | 23.2 | 17.1 | 84.5 | 24.7 | 3.06 | <.001 |
| State-Trait Anxiety<br>Inventory General Score | 33.1 | 8.9 | 57.4 | 9.1 | 9.04 | <.001 |
| State-Trait Anxiety<br>Inventory Present Score | 28.9 | 7.1 | 43.5 | 8.5 | 7.71 | <.001 |
| RT for external attention<br>(seconds) | 2.1 | .47 | 2.1 | .43 | .03 | .826 |
| Accuracy for external<br>attention (%) | 94.4 | 5.8 | 94.0 | 5.7 | .07 | .868 |
| Reaction time for direct<br>self-appraisal (seconds) | 1.8 | .31 | 1.8 | .31 | .23 | .816 |
| Reaction time for reflected<br>self-appraisal (seconds) | 1.7 | .30 | 1.9 | .32 | .64 | .068 |
| Proportion of adjectives<br>affirmed during direct self-appraisal (%) | 45.6 | 10.8 | 51.8 | 12.7 | .53 | .008 |
| Proportion of adjectives<br>affirmed during reflected<br>self-appraisal (%) | 42.7 | 12.2 | 47.9 | 14.3 | .40 | .050 |

### Task performance

There was a statistically significant difference in the RT between the three condition types (*F*(1.28 138.43) = 34.01, *P* < .001) across groups. Post hoc testing revealed significantly slower RTs during the external attention condition compared with both the direct (*P* < .001) and reflected self-appraisal conditions (*P* < .001), as well as a small but significantly slower RT for reflected compared with direct self-appraisal (*P* = .019). There was no significant main effect of diagnosis on RT (*F*(1,108) = 1.21, *P* = .274), nor a significant interaction effect between diagnostic status and RT (*F*(1.28 138.43) = 1.64, *P* = .204). Mean accuracy for the external attention task was 94.4% for controls and 94.0% for SAD participants, with no significant difference between groups (*P* = .868; Table 1). SAD participants were more likely to answer “Yes” to the question “Does this word describe you?” (*t*(108) = -2.68, *P* = 0.008). While there were no significant differences between groups in the percentage of “Yes” responses to the question “Would others use this word to describe you?”, it should be noted that this effect was on the boundary of the statistical significance threshold (*t*(108) = -1.98, *P* = 0.05).

### Mapping common and distinct activations across self-appraisal and rest

Across both healthy controls and SAD participants, direct and reflected self-appraisal conditions produced significant and broadly similar activation patterns which encompassed core regions of the DMN (Figure 1A/B and Figure 2A/B, respectively): namely the anterior medial wall cortex (including frontopolar MPFC), posteromedial cortex (incorporating the PCC and precuneus), and left IPL.

**Figure 1.**
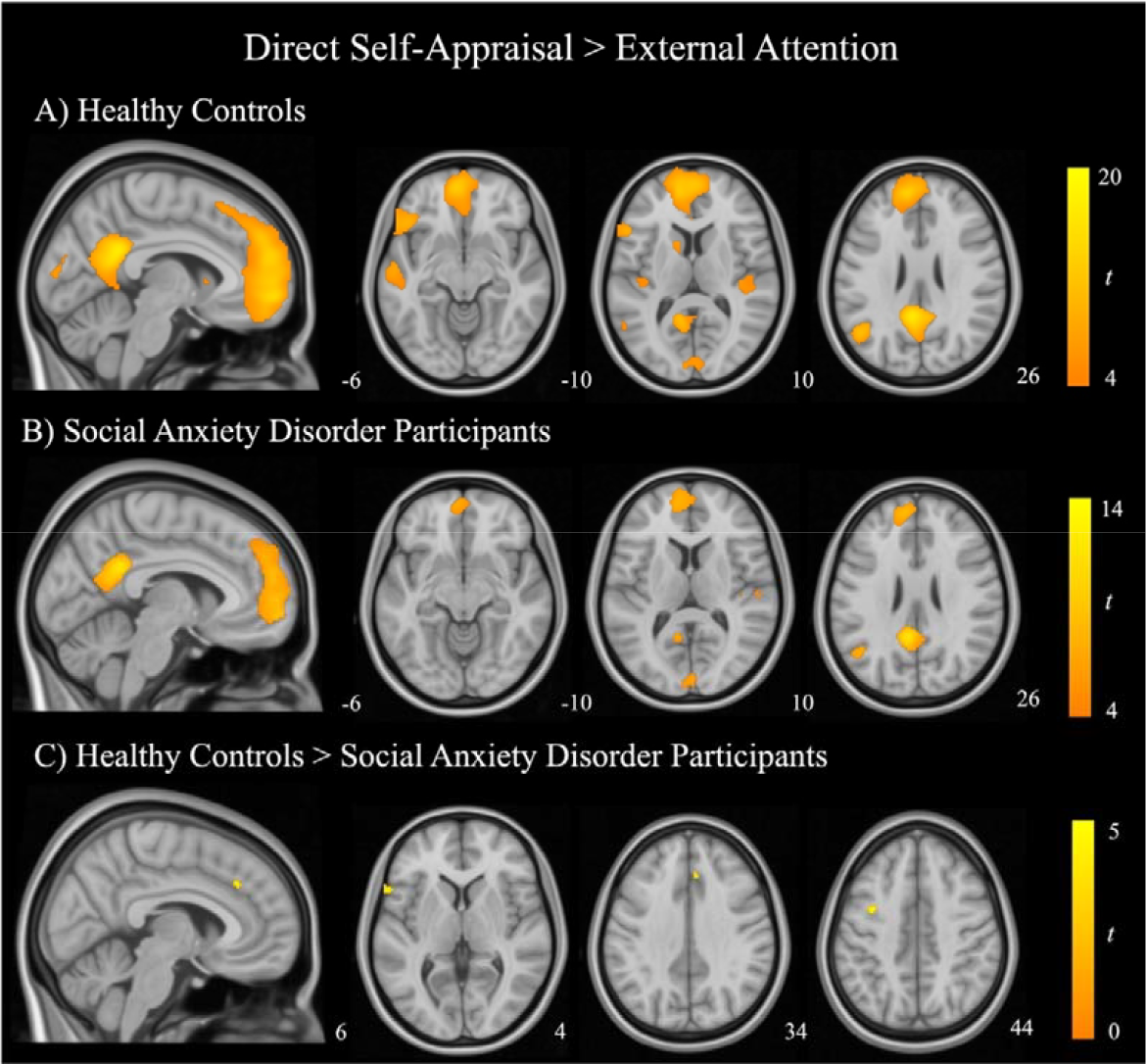
Regions of significant activation during direct self-appraisal compared with external attention for (A) healthy control participants and (B) SAD participants. Results are thresholded at *P*_FWE_ < .05, K_e_ = 20. Also shown is (C) the comparison of healthy controls and SAD participants, which is thresholded at *P* < 0.001, uncorrected, K_e_ = 20. Color bars represent t-statistics.

**Figure 2.**
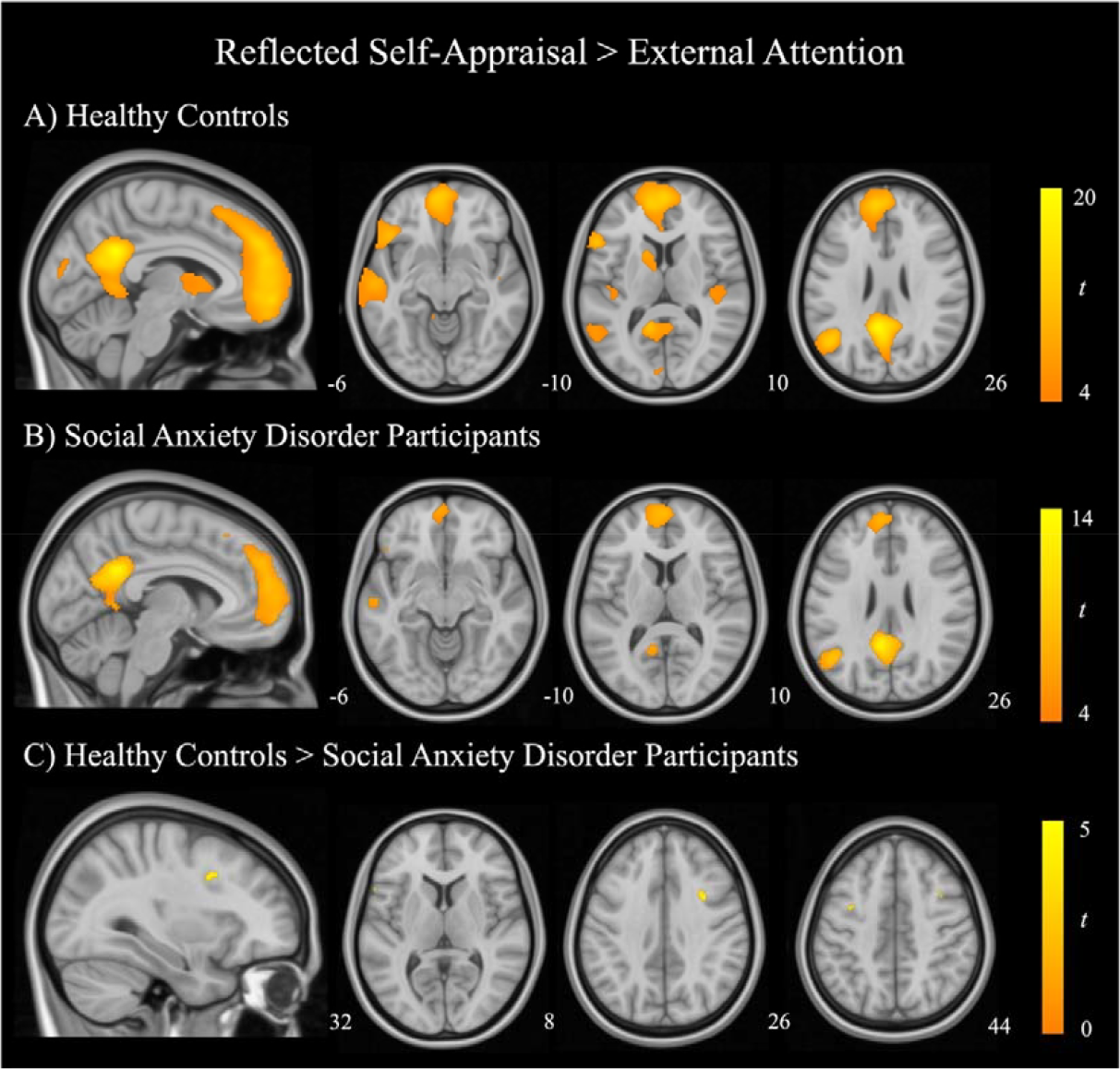
Regions of significant activation during reflected self-appraisal compared with external attention for (A) healthy control participants and (B) SAD participants. Results are thresholded at *P*_FWE_ < .05, K_e_ = 20. Also shown is (C) the comparison of healthy controls and SAD participants, which is thresholded at *P* < 0.001, uncorrected, K_e_ = 20. Color bars represent t-statistics.

The three-way conjunction analysis across participants to identify core-self regions revealed three significant regional clusters: MPFC (peak coordinate, x = -10, y = 54, z = 10; cluster size = 630; peak *t-* value = 7.15), PCC (peak coordinate, x = -6, y = -50, z = 26; cluster size = 158; peak *t*-value = 7.24), and left IPL (peak coordinate, x = -48, y = -60, z = 24; cluster size = 184; peak *t*-value = 6.63; Supplementary Figure S1).

Overall, SAD participants and healthy controls demonstrated similar activation patterns during the direct and reflected self-appraisal conditions. At a whole-brain level, between-groups comparisons revealed no significant differences in regional activation for the main contrasts of interest (*P*_FWE_ < 0.05). At a more lenient threshold (*P* < 0.001, uncorrected), differences were observed for both the direct self-appraisal > external attention and reflected self-appraisal > external attention contrasts.

During direct self-appraisal, SAD participants demonstrated decreased activation in the left pars triangularis of the inferior frontal gyrus, left premotor cortex, and ventral frontal eye fields compared with healthy control participants (Figure 1C; Supplementary Table S2). During reflected self-appraisal SAD participants illustrated decreased activity in the right pars opercularis of the inferior frontal gyrus, left pars triangularis of the inferior frontal gyrus, and left premotor cortex (Figure 2C; Supplementary Table S2).

### Connectivity differences between SAD participants and healthy controls

SAD participants had reduced intrinsic excitatory connectivity from the PCC to MPFC (expected value = -.09 Hz, PP = 1.00) and reduced intrinsic inhibitory connectivity from the left IPL to MPFC (expected value = .08 Hz, PP = 1.00) in comparison to healthy control participants (Figure 3A).

**Figure 3.**
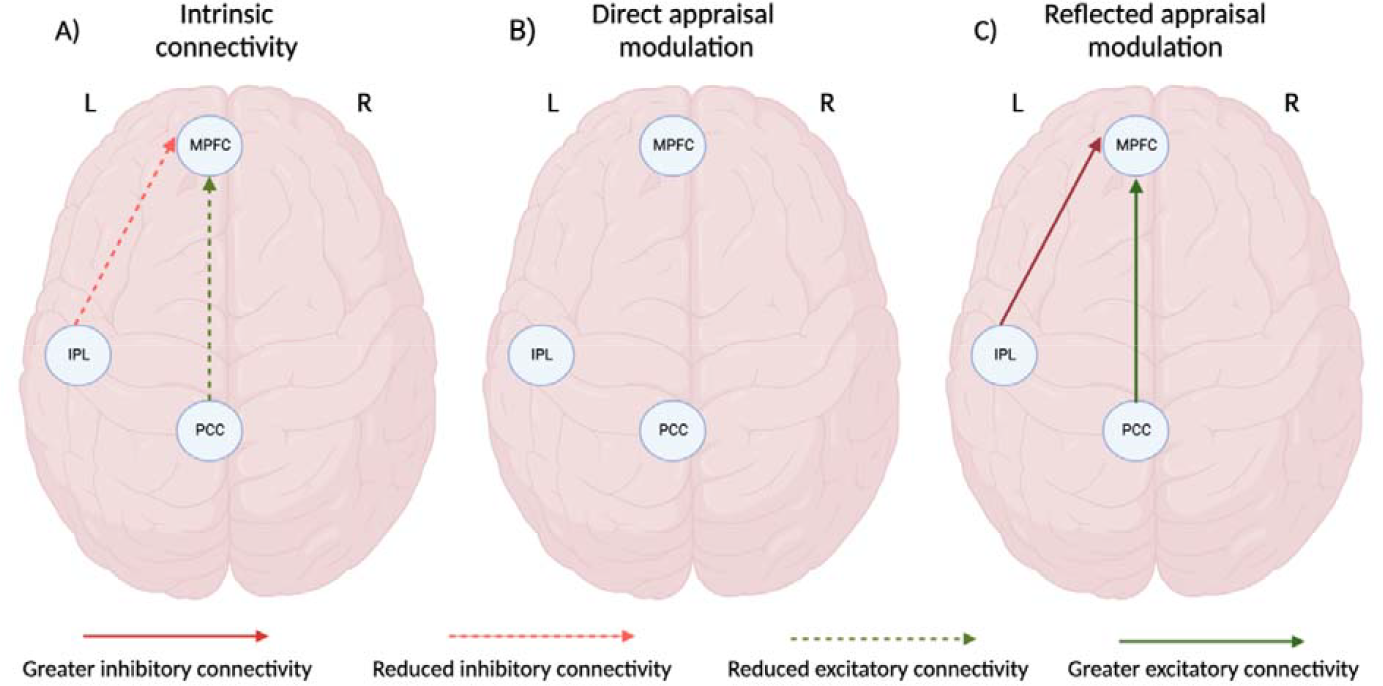
Differences between SAD participants and health controls in the intrinsic connectivity and modulation associated with direct and reflected self-appraisal (PP > .95). Image created with BioRender (www.biorender.com).

During reflected self-appraisal, SAD participants had greater excitatory connectivity from the PCC to MPFC (expected value = .05 Hz, PP = .95) and greater inhibition from the left IPL to MPFC (expected value = -.06 Hz, PP = .97) in comparison with healthy control participants (Figure 3C). No such differences between groups were observed for direct self-appraisal modulation. The expected values and PP for all parameter estimates are reported in Supplementary Table S3-5.

### Leave-one-out cross-validation

Using those parameters which demonstrated between group differences, we performed a LOOCV analysis in the PEB framework to determine whether the size of this effect on these parameters could significantly predict anxiety symptom severity. The intrinsic connectivity from the left IPL to MPFC and from the PCC to MPFC resulted in a significant out-of-samples correlation between the predicted and observed STAI general scores (*r* = .25, *P* = .004; Figure 4A). No significant correlations were found for these intrinsic parameters and LSAS or STAI present scores (Figure 4B and C). For the full list of individual parameter correlations see Supplementary Table S6.

**Figure 4.**
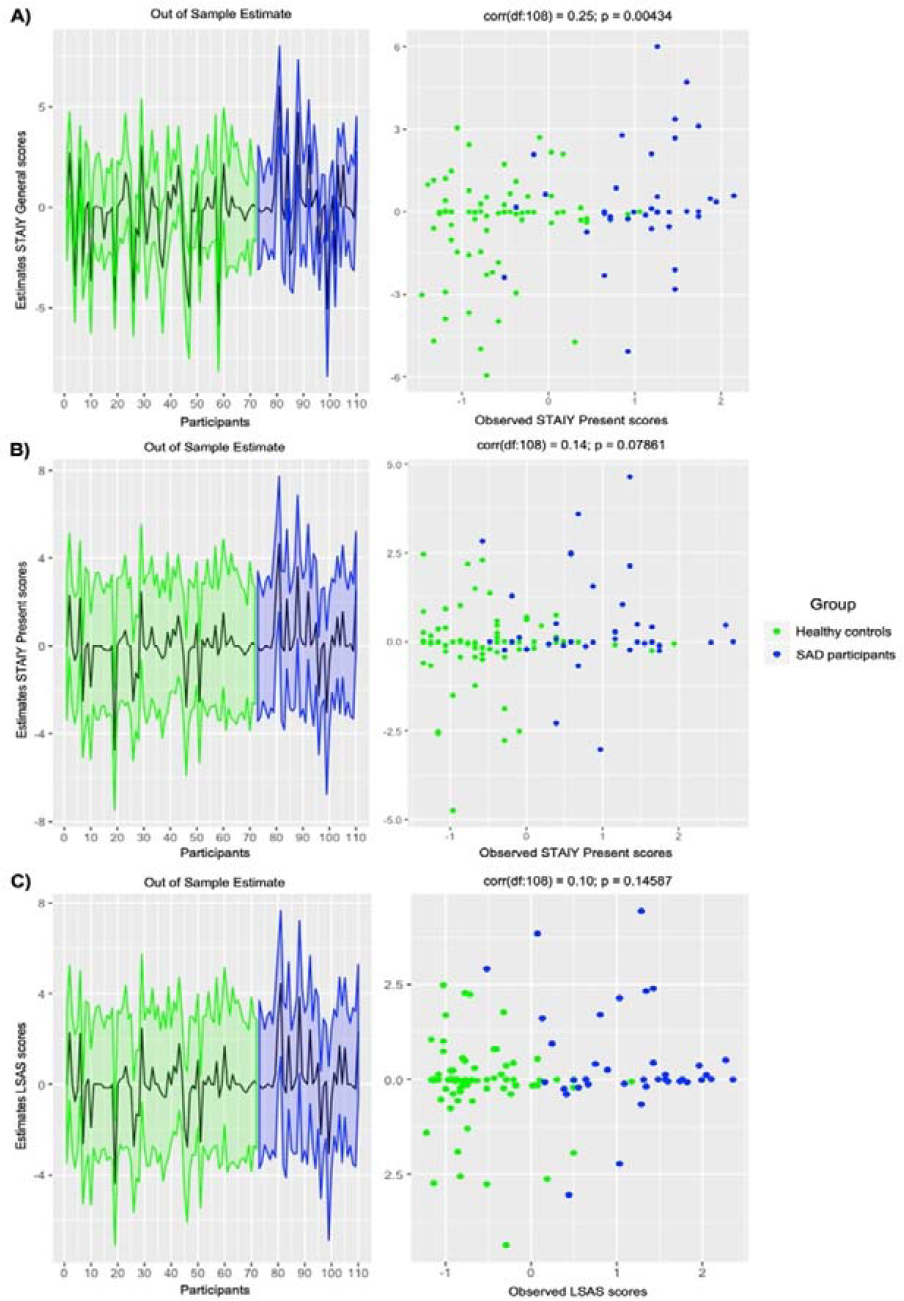
Leave-one-out cross-validation predicting anxiety symptom severity across all participants. **Left:** The out-of-sample estimate of anxiety symptom severity (standardized mean-centered) with 90% credible interval (shaded area). **Right:** The correlation between observed scores and the predicted values for the (A) State-Trait Anxiety Inventory General Subscale, (B) State-Trait Anxiety Inventory Present Subscale, and (C) Liebowitz Social Anxiety Scale across all participants. Data from healthy controls are shown in green; data from SAD participants are shown in blue.

## Discussion

The current study investigated a brain model of the network dynamics present during self-referential processing in young people with SAD. We observed DMN connectivity differences in SAD participants compared with healthy controls during reflected, but not direct self-appraisal. Specifically, during reflected self-appraisal SAD participants illustrated greater excitatory connectivity from the PCC to MPFC and greater inhibitory connectivity from the left IPL to MPFC. The specificity of this finding for the reflected self-appraisal condition is consistent with the nature of SAD, whereby those with SAD show acute, anxiety-inducing preoccupation with the views that others hold of them.

The PCC has been highlighted as a key region in coordinating the function of the DMN during social processes^47^. Our previous work hypothesized that representations of the narrative self, the aspect underlying direct and reflected self-appraisal, are primarily generated by the PCC^40^. In contrast to direct self-appraisal, however, reflected self-appraisal appears to be marked by greater regulatory feedback from the left IPL due to its greater dependence on perspective taking and episodic memory retrieval^31^. The role of the MPFC has been suggested to gate these representations of the self into conscious awareness^15, 48^ and to be more broadly involved in evaluative processes^49, 50^.

While SAD participants have demonstrated both hyper- and hypoconnectivity of the DMN across studies, more recent work has shown reduced resting-state connectivity within the DMN^51, 52^. Broadly, these observations and our own are consistent with the Topography of the Anxious Self model proposed by Angeletti and colleagues^53^. In short, this model suggests that abnormal resting-state connectivity within the DMN and between the DMN and salience network may contribute to trait features of anxiety; these baseline alterations, in turn, increase the sensitivity to task associated hyperactivity of these same regions and increases the likelihood that specific stimuli induce state anxiety. This interpretation is additionally supported by our intrinsic connectivity parameters predicting trait anxiety symptoms. Interestingly, we did not observe effective connectivity alterations during direct self-appraisal in this sample, in contrast to previous findings^32^. Notably, however, in the research by Davey and colleagues^32^, this effect was identified in participants with any comorbid anxiety disorder, not just social anxiety. Therefore, it would be expected that alterations to resting-state connectivity may be consistent across anxiety disorders whereas task related modulation may be more specific to the disorder under consideration. Examining a larger and more heterogeneous sample will allow for direct testing of this hypothesis.

Cognitive models of social anxiety have proposed that increased self-focused attention is a core feature in the development and maintenance of this disorder^54^. A heightened level of self-focused attention has been shown to correlate with the activity of the MPFC and PCC in participants who are highly socially anxious^55^. During reflected self-appraisal, the increased modulation from the left IPL and PCC to MPFC may indicate a heightened focus on how the self is perceived to be represented by others in those with SAD. Interestingly, in SAD participants the connectivity from the PCC to MPFC during reflected self-appraisal also appears more similar to the modulation present during direct self-appraisal. This supports the idea that for SAD participants reflected self-appraisal is more dependent on their constructed narrative self, in contrast to how this process functions in healthy participants^31^. Notably, CBT has been shown to reduce self-focused negative thoughts and beliefs^13^, with decreased frequency of negatively biased thinking being associated with reduced social anxiety^56^. Thus, the activity and connectivity of DMN regions may provide insight into which participants will respond optimally to psychotherapy. Previous research has demonstrated that anterior components of the DMN, including the ACC and MPFC, are associated with treatment response for those with anxiety disorders. Pretreatment activity of the ACC, both at rest^57^ and during an implicit regulation task^58,^ has been associated with improved outcomes to CBT.. Work by Yuan et al^59^ has additionally shown that baseline connectivity between the dorsal MPFC and amygdala are both predictive of response to CBT and is normalized following treatment. As such, examining whether DMN effective connectivity is predictive of treatment response to CBT, and whether successful treatment normalizes this altered connectivity, may be of interest for future research.

This study provides insight into SAD associated alterations to the directional interactions of the DMN, however, there are several limitations which should be considered. The block design of the task and the selection of trait adjectives distributed around the median rating for ‘likeableness’ meant that we were unable to disentangle if the observed effects may have been different for positive and negatively valence adjectives. While recent work has illustrated no significant differences in DMN activity between positive and negative direct self-referential processing^60^, there may be differences present during reflected self-referential processing. While the intrinsic connectivity generated during this task has been compared to findings from resting-state studies, these forms of connectivity are not directly comparable. This highlights that future studies should directly investigate resting state effective connectivity in SAD participants to examine whether these effects are observed in “true” resting states. The SAD sample used in this study was also relatively small, which makes replication in a larger sample of adolescents and young adults with SAD critical.

## Conclusions

Our study has provided a novel brain model of DMN dysfunction present in SAD participants during self-referential processing. We observed SAD associated reductions in the intrinsic connectivity to the MPFC as well as greater modulation to the MPFC specifically during reflected self-referential processing. These observations highlight anxiety associated alterations to connectivity which occur during reflected self-appraisal. We also demonstrated that the effect size of these alterations in the intrinsic connectivity were associated with trait anxiety symptoms. Further investigations into how psychological treatments which specifically target this appraisal process, such as CBT, alters this dysfunction may provide key insight into the neurobiology of SAD.

## CRediT Authorship Contribution Statement

AJJ: Conceptualization; Formal analysis; Methodology; Writing - original draft; Visualization. BJH: Conceptualization, Funding acquisition; Methodology, Supervision; Writing - review & editing. RD: Formal analysis; Writing - review & editing. LS: Resources; Writing - review & editing. KLF: Resources; Writing - review & editing. LP: Resources; Writing - review & editing. CGD: Conceptualization; Funding acquisition; Methodology; Supervision; Writing - review & editing.

## Acknowledgments

The authors thank Dr. Laura Finlayson-Short, Dr. Yara Toenders, Dr. Hannah Savage and Ms. Lisa Incerti for their contributions to data collection. We also thank the staff from the Sunshine Hospital Medical Imaging Department (Western Health, Melbourne).

