## Supplementary Materials for "A brain model of altered self-appraisal in social anxiety disorder"

**Supplementary Methods……………………………………………………………………2**

**Supplementary Tables………………………………………………………………………3**

Supplementary Table S1: Word Lists for the Direct Self-Appraisal, Reflected Self-Appraisal, and External Attention Conditions…………………………………….…………………..….3

Supplementary Table S2: Significantly Greater Activation in Healthy Controls Compared with Social Anxiety Disorder Participants during Direct and Reflected Self-Appraisal…......4

Supplementary Table S3: *Parametric Empirical Bayes Estimates for Intrinsic Connectivity Parameters and their Posterior Probabilities*……..………………………………………….5

Supplementary Table S4: *Parametric Empirical Bayes Estimates for Direct Self-Appraisal Connectivity Parameters and their Posterior Probabilities*…..…………………………..….6

Supplementary Table S5: *Parametric Empirical Bayes Estimates for Reflected Self-Appraisal* *Connectivity Parameters and their Posterior Probabilities*……………………………..…...7

Supplementary Table S6 : *Out-of-samples Correlation Between Anxiety Symptom Severity Using Leave-one-out Cross-validation and the Parameters of Interest*……………………...8

**Supplementary Figures……………………………………………………………………..9**

*Supplementary Figure S1.* Conjunction analysis identifying regions which are commonly activated for rest, direct and reflected self-appraisal conditions compared with the external attention condition, but which illustrated greater activation during direct and reflected self-appraisal compared with rest.………..……………………………….…..…………………..9

Supplementary Figure S2: Model of intrinsic connections (black) and direct input (grey) specified in our DCM analysis ……………………………………………………….….….10

*Supplementary Figure S3.* Distribution of (A) Liebowitz Social Anxiety Scale scores, (B) State-Trait Inventory General and (C) Present scores by diagnostic group…………………11

**Supplementary Methods**

**Propensity score matching**

Propensity score matching was conducted using the R package *MatchIT* with the variable ratio "extremal" matching as described by Ming and Rosenbaum ^1^ and the optimal distance method ^2^. Rather than requiring all patients to be matched to a single control, variable ratio matching allows the number of matches for each patient to vary (which we have limited to between one and two controls for this analysis). This was selected as matching with a variable number of controls has been suggested to increase the precision of matching ^3^, and result in greater reductions to the biasing effect of covariates compared with using a fixed number ^1, 4^. The optimal distance method focuses on minimizing the average absolute distance across all matched pairs. Participants were matched on both age and gender.

### Statistical analysis

Between-group differences in clinical and behavioral characteristics were compared using SPSS version 27 (IBMCorp., Armonk, NY). Reaction times (RT) between groups were compared using a repeated measures ANOVA; due to violating assumptions of sphericity this was estimated using the Greenhouse-Geisser correction. Accuracy differences in the external attention condition were compared with Mann-Whitney U tests. Analyses were adjusted for multiple comparisons using the Benjamini–Hochberg correction ^5^ to determine significance (*P* < 0.05). Previous estimates using frequentist statistics in DCM have suggested that at least 27 participants are required in each group to detected an effect with a Cohen’s *d* of .03 at 80% power ^6^. This information was used to guide the minimum sample size required for this study.

**Supplementary Tables**

Supplementary Table S1

*Word Lists for the Direct Self-Appraisal, Reflected Self-Appraisal, and External Attention Conditions*

| List A | List B | List C |
| --- | --- | --- |
| excited  absent-minded  unintelligent  noisy  unhealthy  argumentative  proud  conformist  moody  rebellious  persuasive  studious  silent  daring  impulsive  cautious  orderly  lonesome  shy  indecisive  self-confident  unobservant  practical  inconsistent  outgoing  clumsy  superstitious  restless  self-assured  impractical | perfectionistic  persistent  unconventional  dependent  inquisitive  unhappy  depressed  sarcastic  confident  overconfident  unpunctual  sociable  calm  lonely  insecure  materialistic  critical  idealistic  careful  thrifty  domineering  emotional  self-critical  short-tempered  gullible  forgetful  daydreamer  curious  oversensitive  unimaginative | serious  inattentive  aggressive  unsociable  nervous  fearful  unpredictable  nonconforming  excitable  untidy  unemotional  indifferent  systematic  obedient  relaxed  timid  wasteful  quiet  talkative  self-conscious  possessive  modest  sentimental  bashful  angry  pessimistic  easy going  bold  tidy  stubborn |

The lists of personality adjectives were drawn from Anderson et al. (1968). They were matched on valence and number of vowels, and the list that comprised each condition was counterbalanced between participants.

Supplementary Table S2

*Significantly Greater Activation in Healthy Controls Compared with Social Anxiety Disorder Participants during Direct and Reflected Self-Appraisal*

| Brain region | BA | Coordinates | | | Cluster size (2mm^3^ voxels) | *t*-value |
| --- | --- | --- | --- | --- | --- | --- |
|  |  | X | Y | Z |  |  |
| *Direct Self-Appraisal > External Attention* | | | | | |  |
| Pars triangularis | 45 | -62 | 18 | 4 | 91 | 4.12 |
| Premotor cortex | 6 | -36 | 0 | 44 | 61 | 4.12 |
| Frontal eye fields/dACC | 8/32 | 6 | 30 | 34 | 44 | 3.59 |
| *Reflected Self-Appraisal > External Attention* | | | | | |  |
| Pars opercularis | 44 | 32 | 8 | 26 | 115 | 4.07 |
| Premotor cortex | 6 | -36 | 2 | 44 | 63 | 3.67 |
| Pars triangularis | 45 | -56 | 18 | 8 | 23 | 3.46 |

Supplementary Table S3

*Parametric Empirical Bayes Estimates for Intrinsic Connectivity Parameters and their Posterior Probabilities*

|  | Mean Connectivity | | Group Difference  (SAD > Controls) | | Age | |
| --- | --- | --- | --- | --- | --- | --- |
| Intrinsic Connectivity | Expected Value | PP | Expected Value | PP | Expected Value | PP |
| MPFC→MPFC | **-.16** | **1.00** | .00 | .57 | .00 | .58 |
| MPFC→PCC | ***-.41*** | ***1.00*** | -.05 | .83 | .00 | .55 |
| MPFC→PCC | ***-.18*** | ***1.00*** | .00 | .73 | .00 | .75 |
| PCC→MPFC | ***.38*** | ***1.00*** | ***-.09*** | ***1.00*** | ***.02*** | ***1.00*** |
| PCC→PCC | ***-.52*** | ***1.00*** | .00 | .51 | ***-.06*** | ***1.00*** |
| PCC→IPL | ***.50*** | ***1.00*** | -.36 | .92 | .00 | .61 |
| IPL→MPFC | ***-.16*** | ***1.00*** | ***.08*** | ***1.00*** | ***-.03*** | ***1.00*** |
| IPL→PCC | ***-.33*** | ***1.00*** | .00 | .54 | -.02 | .85 |
| IPL→IPL | .00 | .55 | .00 | .62 | ***.04*** | ***1.00*** |

Supplementary Table S4

*Parametric Empirical Bayes Estimates for Direct Self-Appraisal Connectivity Parameters and their Posterior Probabilities*

|  | Mean Connectivity | | Group Difference  (SAD > Controls) | | Age | |
| --- | --- | --- | --- | --- | --- | --- |
| Direct Self-Appraisal Connectivity | Expected Value | PP | Expected Value | PP | Expected Value | PP |
| MPFC→PCC | ***-.37*** | ***1.00*** | -.01 | .60 | .00 | .51 |
| MPFC→IPL | -.01 | .57 | .00 | .50 | .00 | .60 |
| PCC→MPFC | ***.31*** | ***1.00*** | .01 | .59 | -.02 | .90 |
| PCC→IPL | ***.18*** | ***1.00*** | .01 | .62 | .01 | .71 |
| IPL→MPFC | -.01 | .59 | -.02 | .75 | ***.03*** | ***.95*** |
| IPL→PCC | .01 | .54 | -.06 | .90 | .00 | .56 |

Supplementary Table S5

*Parametric Empirical Bayes Estimates for Reflected Self-Appraisal* *Connectivity Parameters and their Posterior Probabilities*

|  | Mean Connectivity | | Group Difference  (SAD > Controls) | | Age | |
| --- | --- | --- | --- | --- | --- | --- |
| Reflected Self-Appraisal Connectivity | Expected value | PP | Expected value | PP | Expected value | PP |
| MPFC→PCC | ***-.90*** | ***1.00*** | .04 | .73 | .01 | .68 |
| MPFC→IPL | ***-.06*** | ***.98*** | .02 | .72 | .01 | .80 |
| PCC→MPFC | ***.25*** | ***1.00*** | ***.05*** | ***.95*** | .01 | .72 |
| PCC→IPL | ***.25*** | ***1.00*** | -.01 | .64 | -.01 | .77 |
| IPL→MPFC | -.01 | .67 | ***-.06*** | ***.97*** | -.00 | .56 |
| IPL→PCC | ***-.47*** | ***1.00*** | .00 | .52 | .00 | .57 |

Supplementary Table S6

*Out-of-samples Correlation Between Anxiety Symptom Severity Using Leave-one-out Cross-validation and the Parameters of Interest*

| Parameters | Correlation | *P*-value |
| --- | --- | --- |
| LSAS |  |  |
| *Intrinsic Connectivity* |  |  |
| IPL → MPFC | .05 | .31 |
| PCC→ MPFC | .09 | .18 |
| *Reflected Self-appraisal Modulation* |  |  |
| IPL → MPFC | -.04 | .68 |
| PCC→ MPFC | -.53 | 1.00 |
| STAI general |  |  |
| *Intrinsic Connectivity* |  |  |
| IPL → MPFC | .17* | .04 |
| PCC→ MPFC | .15 | .05 |
| *Reflected Self-appraisal Modulation* |  |  |
| IPL → MPFC | .06 | .25 |
| PCC→ MPFC | -.05 | .69 |
| STAI present |  |  |
| *Intrinsic Connectivity* |  |  |
| IPL → MPFC | .02 | .40 |
| PCC→ MPFC | .14 | .07 |
| *Reflected Self-appraisal Modulation* |  |  |
| IPL → MPFC | -.02 | .58 |
| PCC→ MPFC | -.38 | 1.00 |

*Note.* * Significant at *p* < .05

**Supplementary Figures**

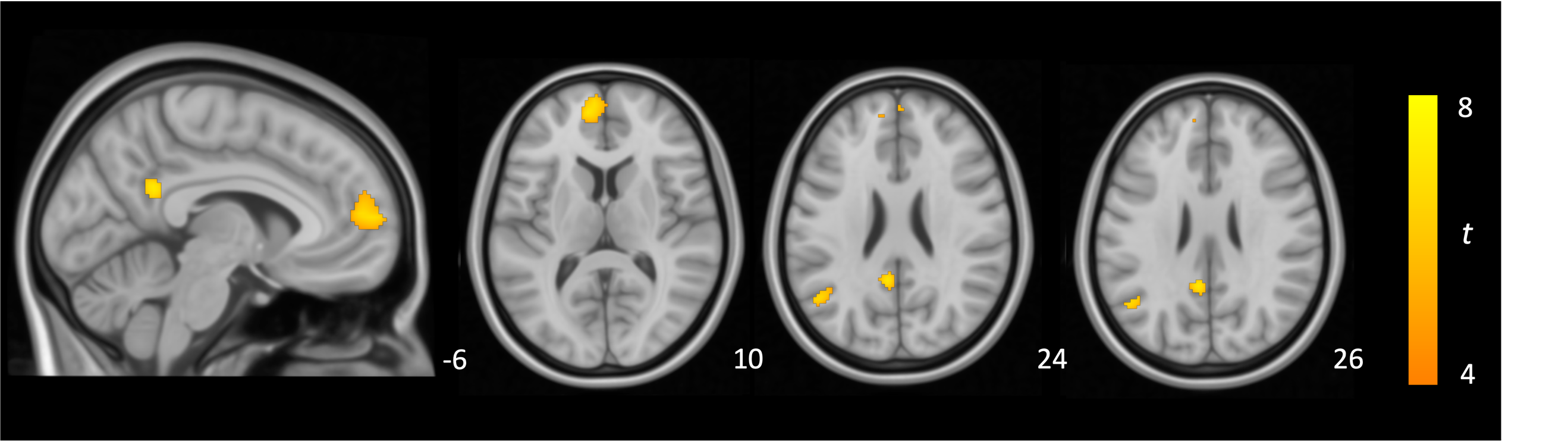

*Supplementary Figure S1.* Conjunction analysis identifying regions which are commonly activated for rest, direct and reflected self-appraisal conditions compared with the external attention condition, but which illustrated greater activation during direct and reflected self-appraisal compared with rest. Results thresholded at *P*_FWE_ < 0.05. Ke = 20. Color bar represents t-statistic.

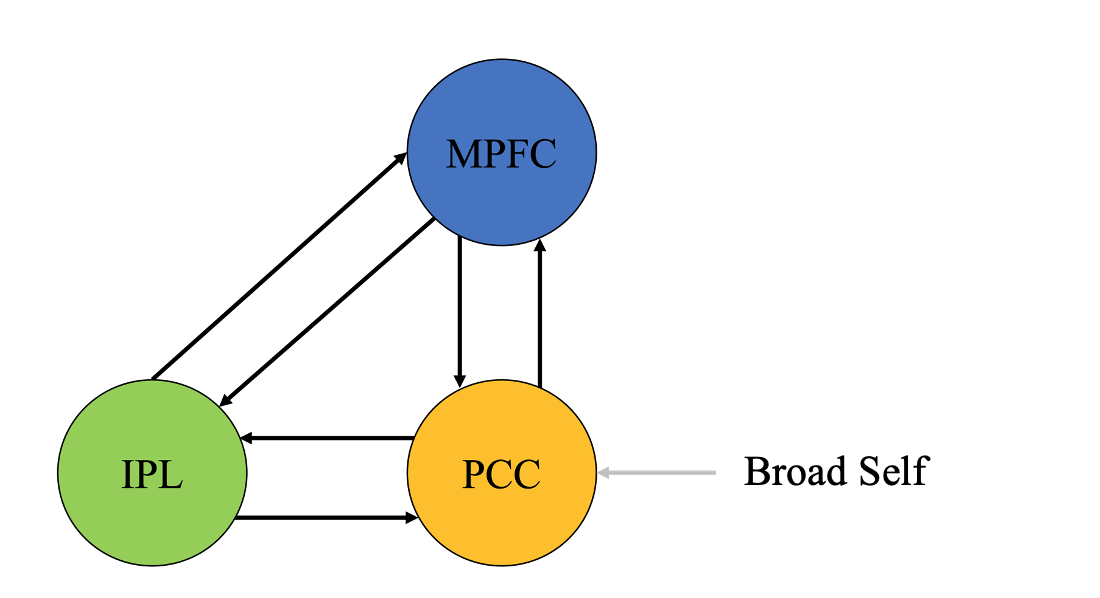

*Supplementary Figure S2.* Model of intrinsic connections (black) and direct input (grey) specified in our DCM analysis.

*Note.* MPFC, medial prefrontal cortex; PCC, posterior cingulate cortex; IPL, inferior parietal lobule.

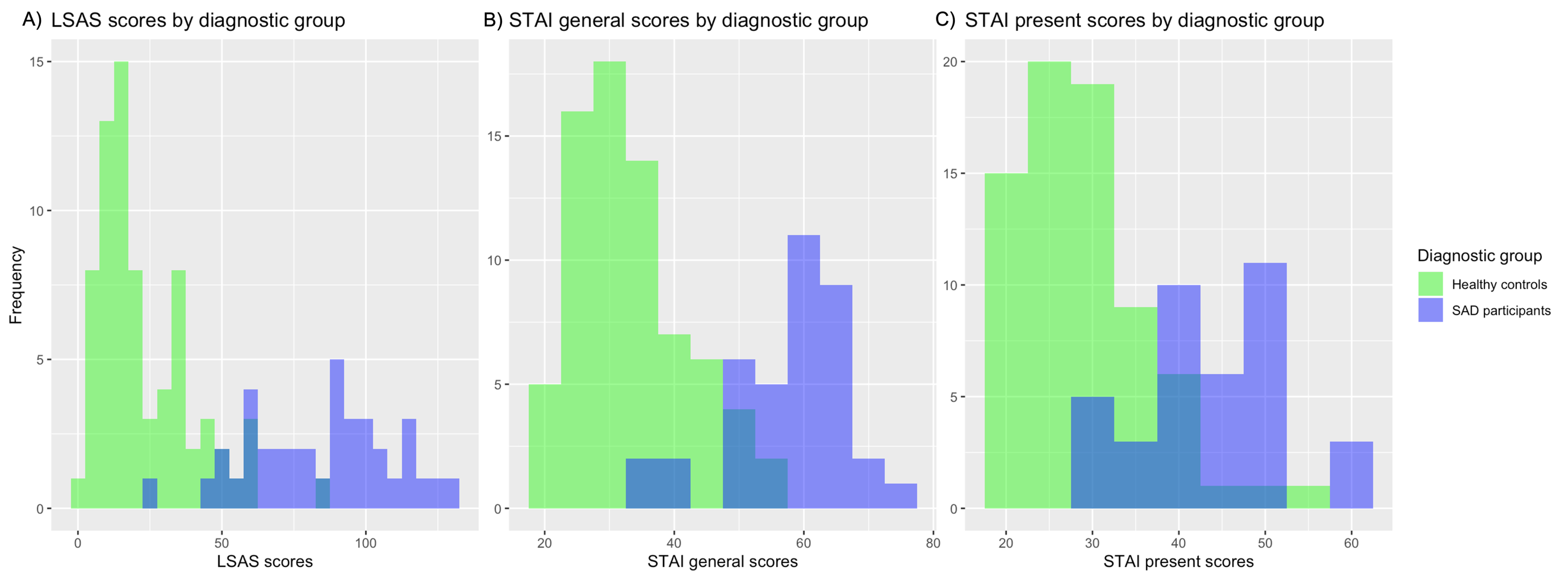
 *Supplementary Figure S3.* Distribution of (A) Liebowitz Social Anxiety Scale scores, (B) State-Trait Inventory General and (C) Present scores by diagnostic group.
